# Parental B-MYB/FOXM1 controls mitotic E2F to determine daughter cell fate

**DOI:** 10.64898/2026.09.21.753170

**Authors:** David L. Rosenthal, Tatsuki Tsuruoka, Yumi Konagaya, Bo Gong, Vinodhini Govindaraj, Samsara W. Upadhya, Nalin Ratnayeke, Debarya Saha, Wenxin Xia, Mary N. Teruel, Tobias Meyer

**Affiliations:** Department of Biochemistry and Biophysics, Weill Cornell Medicine, New York, NY, USA; Drukier Institute for Children’s Health, Weill Cornell Medicine, New York, NY, USA

## Abstract

Mitogens trigger cell-cycle entry by activating E2F at the restriction point, which is followed by B-MYB/FOXM1 activation and progression to mitosis. How mitogens control continued cycling and cell-cycle exit after the restriction point is not well-understood. By developing an E2F and B-MYB/FOXM1 dual transcriptional biosensor system, we show that S/G2 phase duration is set by timed mitogen-regulated B-MYB/FOXM1 activation, while E2F activity gradually declines before mitosis. As a striking consequence, rapid B-MYB/FOXM1 activation shortens S/G2, delivering high mitotic E2F activity to daughter cells which keeps them cycling. Delayed B-MYB/FOXM1 activation prolongs S/G2, depleting mitotic E2F which drives daughters to quiescence. When S/G2 is further prolonged, partially activated B-MYB/FOXM1 frequently reverts, triggering mitotic bypass and polyploid quiescence. Thus, B-MYB/FOXM1 governs a tri-directional “G2 restriction point” where cells commit to continued cycling through early B-MYB/FOXM1 activation; cell-cycle exit through delayed B-MYB/FOXM1 activation; or mitotic bypass by B-MYB/FOXM1 inactivation.

## INTRODUCTION

The classic cell-cycle model states that cells sense mitogens in G1, where a restriction point (R) in late G1 determines not only whether quiescent cells enter the cycle but also cycling cells continue to divide ^1^. After passing this point, cells have been thought to complete S, G2, and mitosis (M) independently of further mitogen stimulation (Fig. 1A). However, two recent observations challenge this model. First, for daughters to keep cycling after mitosis, mitogen stimuli were shown to be required during S and G2 (S/G2) of the mother cell, but were not needed after mitosis in G1 of daughter cells ^2–4^. Second, low mitogen levels or CDK4/6 inhibition, that were both believed to block proliferation by suppressing E2F activity in G1, were unexpectedly found to promote mitotic bypass and polyploid quiescence when applied during the mother cell’s S/G2 ^5–7^. To align the cell-cycle model with these new findings, a mechanistic understanding is needed of how cells in S/G2 control the three observed fates: continued cycling, cell-cycle exit to quiescence after mitosis, and mitotic bypass to polyploid quiescence (Fig. 1B).

**Figure 1:**
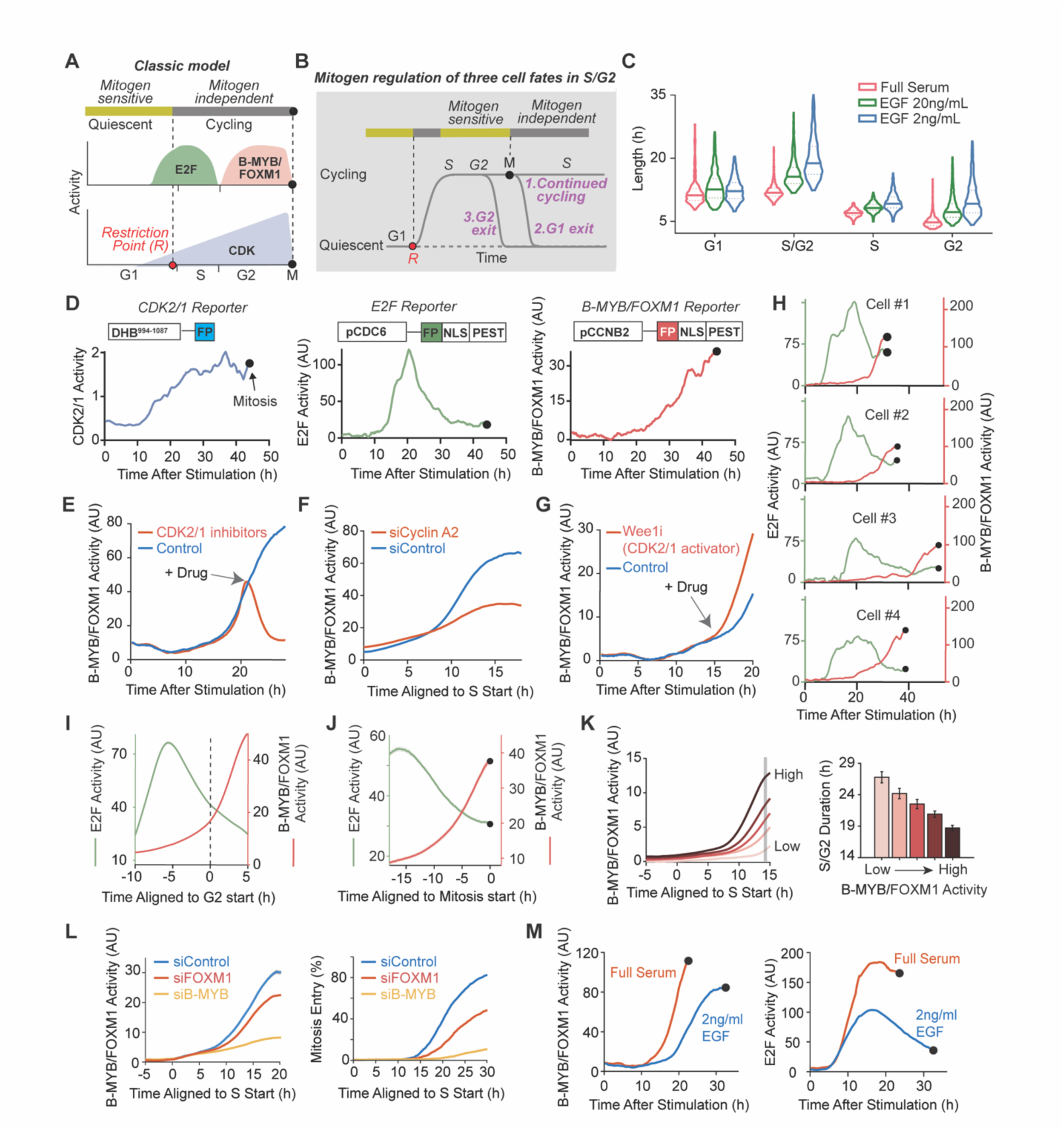
Mitogen stimuli accelerate B-MYB/FOXM1 activation to shorten S/G2 length. **(A)** Classic cell-cycle model: canonical mitogen sensitivity and restriction point, E2F and FOXM1 transcriptional waves, and CDK2/1 activation ramp. (**B**) Mitogens in S/G2 control three cell-cycle fates, which are inconsistent with the classic model. (**C)** Violin plot showing the probability density of cell-cycle phase durations for cells stimulated either by full-serum media (n = 1010), 20 ng/mL EGF (n = 2363), or 2 ng/mL EGF (n = 1343) (measured by CRL4^Cdt2^ degron reporter). (**D)** Schematics of three live-cell reporter activity traces from a single cell after stimulation with 2 ng/mL EGF. Black dots mark mitosis (2 h before anaphase). **(E)** Mean FOXM1 activity, comparison of control (n = 4028) and combined addition of CDK1 inhibitor (CDK1i, Ro-3306 10μM) and CDK2 inhibitor (CDK2i, PF-07104091 1μM) (n = 3690) added at 20 h after stimulation (**F**) Mean FOXM1 activity, comparing treatments with siCCNA2 (Cyclin A2) (n = 3082) or siControl (n = 2946). (**G)** Mean FOXM1 activity time-courses comparing cells treated at 15 h (arrow) with 110 nM MK-1775 (Wee1i; n = 1456) or DMSO (n = 1190). **(H)** Examples of simultaneous single-cell E2F and B-MYB/FOXM1 activity traces (2 ng/mL EGF) (**I)** Mean E2F and B-MYB/FOXM1 activities, computationally aligned to G2 start (defined using CRL4^Cdt2^ reporter); (FOXM1, n = 3262 cells, E2F, n = 929 cells; +/-SEM, shaded area mostly hidden behind mean line). (**J)** Mean E2F and B-MYB/FOXM1 activities (+/-SEM) (n = 1943); computationally aligned to mitosis (1 h before anaphase). (**K**) Left: Mean B-MYB/FOXM1 activity traces of cells computationally sorted into five bins by relative B-MYB/FOXM1 activity 14 h after S start (defined by CDK2/1 reporter). Right: mean S/G2 length +/-2*SEM as a function of relative B-MYB/FOXM1 activity (n = 1468). (**L)** Comparison of siControl, siB-MYB, and siFOXM1 conditions, mean B-MYB/FOXM1 activity (left) and % of cells entering mitosis (right) over time since S start. 2 ng/mL EGF (siControl, n = 1722; siB-MYB, n = 1811; siFOXM1, n = 1610). (**M**) Mean B-MYB/FOXM1 and E2F activity time courses for cells completing mitosis, comparing full-serum media (n = 2298) and 2 ng/mL EGF (n = 2834). All experiments repeated >= 2 times on different days unless noted. Each condition evaluated in 2-4 different wells in 96-well plates.

What has been established is that quiescent cells pass the restriction point by activating E2F transcription factors (E2F1/2/3) ^8,9^, which promote a steady increase of cyclin-dependent kinase (CDK) activity (a combined CDK2/1 activity) (Fig. 1A) ^4,10^. E2F also induces the expression of the pro-mitotic transcription factors B-MYB (encoded by *MYBL2* gene) and FOXM1 ^11,12^. Although the regulation of E2F in G1 and S phases is well characterized, the transcriptional coordination of the cell cycle after S-phase has received less attention. It has been shown that after cells enter S phase, E2F-induced E2F7/8 inhibits E2F1/2/3 ^13^, together with additional mechanisms that inactivate and degrade E2F1/2/3 (Fig. 1A) ^14,15^. Furthermore, beginning in late S-phase, B-MYB and FOXM1 bind to the MuvB transcriptional complex and are activated through phosphorylation by CDK2/1 to induce Cyclin A and mitotic proteins ^11,12,16–18^. The transcriptional activity of MuvB in complex with B-MYB and FOXM1, hereafter referred to as “B-MYB/FOXM1 activity”, is required for initiation of mitosis ^19^. At the end of mitosis, APC/C^Cdh1^ is activated to degrade cyclin A2, FOXM1 and E2F7/8 to initiate the subsequent G1 phase (Fig. 1A) ^20,21^. It was previously thought that continued cycling of daughter cells after mitosis required that mitogens again activate E2F in G1 to reach the restriction point. Recent studies have shown that mitogen signaling in S/G2 can control daughter-cell proliferation independent of mitogen sensing in G1, and that mitogen signals in G2 can prevent premature G2 exit ^3,5–7^. These results suggest that core restriction-point control extends beyond G1 and includes the previous S/G2 (Fig. 1B).

This work was motivated by an unexpected observation we made in initial studies: mitogen signals in S/G2 regulate the length of S/G2. To understand the regulation of S/G2 length and its connection to cell-cycle fate, we developed a live reporter system to measure both B-MYB/FOXM1 and E2F activity changes in S and G2. We show that mitogens regulate the activation of the B-MYB/FOXM1 transcriptional program, thereby controlling the length of S/G2. Contrary to previous work suggesting that E2F is completely inactivated before mitosis ^22^, we found that E2F activity persists until mitosis, though gradually declining from early S onward. Thus, by regulating S/G2 length, the timing of B-MYB/FOXM1 activation controls mitotic E2F activity. Strikingly, a short S/G2 results in high E2F activity being carried through mitosis, which results in daughter cells bypassing the G1 restriction point and continuing to cycle. In contrast, longer S/G2 durations result in low mitotic E2F activity, directing daughters to enter quiescence (G1 exit). Even longer S/G2 durations allow for the reversal of B-MYB/FOXM1 activation in G2, which triggers cell-cycle exit into a polyploid quiescent state, bypassing mitosis (G2 exit). Thus, we propose that the timing of B-MYB/FOXM1 activation and its inactivation constitute a tri-directional “G2 restriction point” that directs cells to either continue cycling, exit the cell cycle post-mitosis, or undergo mitotic bypass.

## RESULTS

### Increasing Mitogen Stimuli Shortens S/G2 Length

If S and G2 are mitogen-independent ^1,8,23^, S/G2 duration should be insensitive to mitogen strength. We challenged this assumption by precisely measuring S and G2 durations in MCF10A breast epithelial cells, a widely used system for studying human cell proliferation. The cells expressed a FUCCI(CA)-2 CRL4^Cdt2^ degron reporter to mark G1/S and S/G2 transitions (Fig. S1A) ^24^, along with an H2B-iRFP670 nuclear marker and a nuclear translocation–based CDK2/1 activity reporter that tracks the rise and fall of the combined cyclin E/A–CDK2/1 activity across the cell cycle (Fig. S1B) ^4,25,26^.

The durations of the G1, S, and G2 phases were measured for low and high EGF and full serum stimuli by using automated tracking and analysis of thousands of cycling cells. Probability distributions showed that the lengths of all cell-cycle phases were variable, particularly for low EGF stimulation. Importantly, the average duration of the combined S/G2 phase, as well as of S and G2 individually, was longer for low mitogen stimulation (2 ng/mL EGF) compared to high EGF stimuli or media with full serum (Fig. 1C). Thus, mitogen stimulation shortens S/G2 duration.

To investigate how S/G2 signaling regulates the length of S/G2 and the different cell-cycle fates, we developed a tripart fluorescent reporter system to simultaneously measure core cell-cycle signaling activities: E2F, B-MYB/FOXM1, and CDK2/1 (Fig. 1A,1D). The E2F reporter consists of a nuclear-localized fluorescent protein under the control of a strong E2F target promoter (pCDC6), along with a destabilization sequence to ensure that both increases and decreases in E2F activity can be measured ^4,25^. To measure B-MYB/FOXM1 activity, we designed a novel B-MYB/FOXM1 activity reporter. Given that cyclin B2 is a major B-MYB/FOXM1 target, and that the cell-cycle dynamics of Cyclin B2 mRNA closely matches that of other B-MYB/FOXM1 targets ^22^, we constructed the reporter around a cyclin B2 (CCNB2) promoter region centered on a B-MYB and FOXM1 binding site (Fig. S1C, D), again inducing a destabilized, nuclear-targeted fluorescent protein (Fig. 1D) ^27^.

Single-cell validation experiments showed that the B-MYB/FOXM1 reporter signal is correlated with the mRNA copy numbers of previously identified B-MYB/FOXM1 targets, cyclin B1 and PLK1 (Fig. S1E) ^22^. To directly determine whether cyclin A2-CDK2/1 activates B-MYB/FOXM1 in G2 ^16,28^, we acutely co-inhibited CDK1 and CDK2, which robustly suppressed B-MYB/FOXM1 activity (Fig. 1E), as did knockdown of the CDK2/1 activator cyclin A2 (Fig. 1F). Conversely, increasing CDK2/1 activity by inhibiting WEE1 kinase accelerated FOXM1 activation (Fig. 1G) ^29^. These complementary perturbations demonstrate that cyclin A2–CDK2/1 is a principal driver of B-MYB/FOXM1 activation during G2.

### Mitogen Stimuli Accelerate B-MYB/FOXM1 Activation to Shorten S/G2 Length

Live-cell imaging of the tripart reporter cells revealed unexpected dynamics of the single-cell E2F and B-MYB/FOXM1 activities during S/G2. Contrary to the prevailing view that E2F activity is fully inactivated after S-phase entry (Fig. 1A) ^14,30^, individual cells retained variable E2F activity through mitosis (Fig. 1H). B-MYB/FOXM1 behaved unexpectedly as well: instead of rising at a fixed time after S start, as previously proposed, B-MYB/FOXM1 activation occurred after variable delays and with variable kinetics across cells ^28^. Despite this heterogeneity, aligning cells to the S–to-G2 transition revealed that the average rate of B-MYB/FOXM1 increase was consistently faster in G2 than in S (Fig. 1I), consistent with previous observations that DNA replication suppresses CDK2/1 and thereby limits FOXM1 activity in S phase ^28,31,32^.

Given the unexpected cell-to-cell variability in B-MYB/FOXM1 activation, we next asked when B-MYB/FOXM1 activity rises relative to mitosis. Aligning individual B-MYB/FOXM1 traces to mitotic entry showed that average B-MYB/FOXM1 activity increases steadily in the hours leading up to mitosis (Fig. 1J). To determine whether the timing of this activation affects S/G2 duration, we grouped cells by their B-MYB/FOXM1 activity 14 hours after S start, a time point corresponding to mid G2 under low-EGF conditions (Fig. 1C). B-MYB/FOXM1 levels at this time were inversely correlated with overall S/G2 length, establishing a link from the relative B-MYB/FOXM1 activity in G2 to the time required to reach mitosis (Fig. 1K). Furthermore, Both B-MYB and FOXM1 knockdown delayed the increase in B-MYB/FOXM1 reporter and entry into mitosis, demonstrating that B-MYB/FOXM1 indeed regulates S/G2 length and mitotic entry (Fig. 1L). Additionally, we determined whether mitogens regulate B-MYB/FOXM1 activity in S/G2. Low EGF not only prolonged S/G2 (Fig. 1C) but also delayed B-MYB/FOXM1 activation (Fig. 1M, left). Correspondingly, E2F activity peaked at nearly the same time and then began to decline, with a large decrease observed in the Low EGF group (Fig. 1M, right). We therefore conclude that mitogen signals shorten S/G2 through accelerated B-MYB/FOXM1 activation.

### Accelerating S/G2 Promotes High Mitotic E2F Activity, Favoring Continued Cycling of Daughter Cells After Mitosis

We next examined the unexpected observation that E2F remains active at mitosis (Fig. 1H, 1J). Despite cells retaining E2F activity in G2, there was no significant correlation between the relative E2F activity in G2 (defined as 14 hours after S start, same as in Fig. 1K) and S/G2 length (Fig. 2A), in contrast to the strong correlation observed with B-MYB/FOXM1 (Fig. 1K). This was consistent with B-MYB/FOXM1, but not E2F, regulating S/G2 length.

**Fig. 2:**
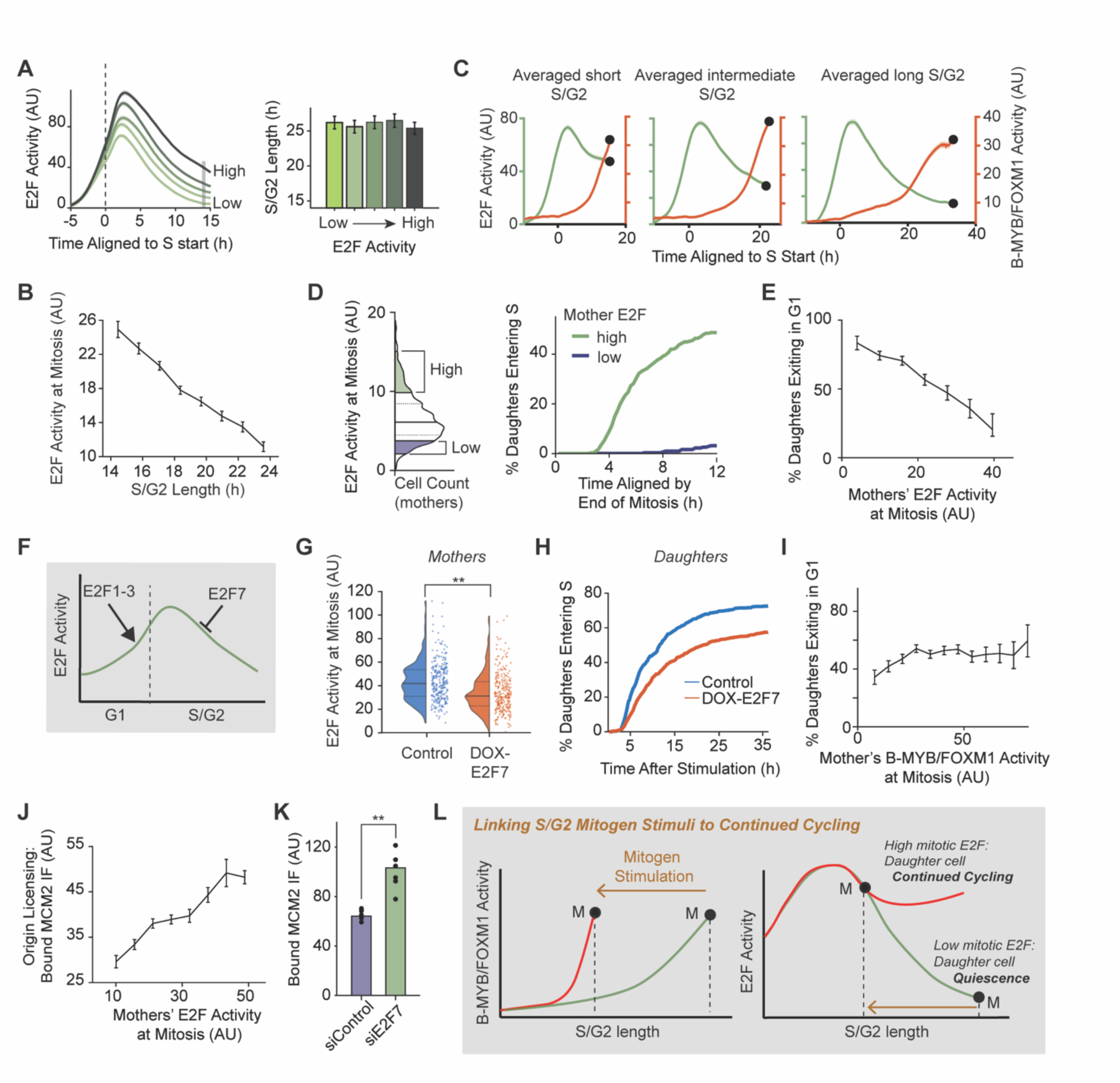
Rapid B-MYB/FOXM1 activation shortens S/G2, Resulting in High Mitotic E2F to Direct Continued Proliferation of Daughters. **(A)** Left: Mean E2F activity traces of cells computationally sorted into five bins based on relative E2F activity 14 h after S entry (defined by CDK2/1 reporter). Right: mean S/G2 length +/-2*SEM of the cells in each relative E2F activity bin (n = 1335). (**B**) Mother cells’ mitotic E2F activity (measured 1-2 hours prior to anaphase) versus S/G2 length, 20 ng/mL EGF stimulation. Computationally sorted into 8 bins based on S/G2 length (n = 2143). (**C)** Mean E2F and B-MYB/FOXM1 activity traces (+/-SEM), 2 ng/mL EGF stimulation; Traces binned by S/G2 length, (Short 1 - 20th percentile; Medium = 40 - 60^th^ percentile; Long = 80 - 99^th^ percentile). (**D)** Left: vertical histogram of E2F activity at mitosis, 20 ng/mL EGF, highlighting “high” and “low” E2F activity cells (n = 1592 cells). Right: Daughter cell cumulative S-phase entry for daughters born of mothers with high vs. low mitotic E2F activity (n = 345 and 314 daughter cells, respectively). (**E)** Percent of daughters undergoing G1 exit plotted as a function of mothers’ E2F activity at mitosis (n = 3257), 7 bins, relative mothers E2F activity. Error bars, 2*SEM. **(F)** Schematic: E2F1/2/3-induced E2F activation in G1 and E2F7/8-mediated E2F suppression in S/G2. (**G)** Violin plot of Mitotic E2F activity; cells expressing Dox-inducible E2F7, treated with doxycycline or DMSO control (n = 300 per condition; difference p = 8.78 * 10^−12^). **(H)** Cumulative S-phase entry in daughter cells of E2F7-overexpressing mother cells (DOX-E2F7, n = 984) or control (DMSO, n = 589). 20 ng/mL EGF; doxycycline or DMSO addition 21 h after release. Analysis limited to cells dividing within 6-10 h of drug addition. **(I)** Same analysis as in (E) but binning by relative mitotic B-MYB/FOXM1 activity at mitosis and plotting the precent of daughter cells exiting to quiescence. (**J)** Chromatin-bound MCM2 immunofluorescence in daughter cells versus E2F activity of their mother at mitosis. Computationally sorted into 8 bins based on mother E2F activity at mitosis (n = 658). Error bars, +/-2*SEM. 20 ng/mL EGF. Cells were gated for addition of 1 μM palbociclib 0-2 h before mitosis to prevent de-novo E2F activation in daughter cells. (**K)** Chromatin bound MCM2 in cells cycling in 20 ng/mL after transfection with siControl (n = 3106 cells) or siE2F7 (n = 2215 cells). Computationally gated for G1 phase. Individual points show well replicates. P = 3.13*10^−4^. (**L)** Schematic of how B-MYB/FOXM1 controls S/G2 length, mitotic E2F activity, and daughter cell fate.

However, E2F activity at mitosis (mitotic E2F activity) had a strong inverse correlation with S/G2 duration (Fig. 2B). Examining E2F activity across S/G2 revealed a slow decline in E2F activity on average from early S phase through mitosis. This slow constant decline is evident when comparing averaged B-MYB/FOXM1 and E2F traces for short, intermediate, and long S/G2 lengths (Fig. 2C). Such analysis reveals that mitotic E2F activity reflects how quickly a cell has transitioned through the previous S and G2 phases. Since cells with high E2F activity are thought to have passed the restriction point ^8,9^, we considered that cells maintaining high E2F through mitosis would not need to pass the restriction point again and would continue cycling unchecked through the subsequent G1/S.

Strikingly, daughters of mothers with high mitotic E2F activity entered the next cell cycle within hours, whereas daughters of mothers with low mitotic E2F almost uniformly exited the cycle (Fig. 2D). Across the full spectrum of mitotic E2F levels, higher mitotic E2F corresponded to fewer daughters exiting to quiescence (Fig. 2E). In line with our hypothesis, the duration of mother-cell S/G2, which inversely correlates with mitotic E2F activity, also inversely correlates with the likelihood of continued daughter-cell cycling (Fig. S2A).

To directly assess whether mitotic E2F activity influences daughter cell-cycle fate, we acutely increased E2F7, a type of repressive E2F, expression using a doxycycline-induced expression system. This method selectively suppresses E2F activity in S/G2 phase, as E2F7 is degraded by APC/C^Cdh1^ in G1 (Fig. 2F) ^33^. Inducing E2F7 lowered mitotic E2F activity (Fig. 2G) and correspondingly reduced the percent of cycling daughter cells (Fig. 2H), demonstrating that mitotic E2F in mother cells controls the continued cycling of daughter cells. We also asked whether high B-MYB/FOXM1 activity at mitosis would similarly reduce the percentage of cells exiting in G1. However, high mitotic B-MYB/FOXM1 levels did not correlate with reduced G1 exit (Fig. 2I), supporting that continued cycling of daughters is driven by high mitotic E2F and not B-MYB/FOMX1.

To continue cycling after mitosis, daughters must rapidly license origins of replication ^34^. We therefore asked whether mitotic E2F activity in mothers controls origin licensing in daughters. After live-cell imaging, we fixed cells and quantified chromatin-bound MCM2 in each daughter (Fig. S2B), then mapped each daughter’s MCM2 level back to its mother’s mitotic E2F activity using RT-QIBC (retrospective time-lapse synchronized quantitative imaged-based cytometry) (Fig. 2J) ^35^. In line with mitotic E2F activity in mothers preparing the daughters for DNA replication, higher mitotic E2F activity in mothers correlated with increased origin licensing in daughters. Moreover, increasing mitotic E2F activity by knocking down E2F7 resulted in higher origin licensing in the daughters (Fig. 2K), demonstrating that mitotic E2F activity controls origin licensing. A time-course analysis further showed that the increase in origin licensing in the daughters caused by knockdown of E2F7 is observable immediately after completion of mitosis (Fig. S2C). Thus, high E2F activity sustained throughout mitosis, by promoting rapid origin licensing, promotes rapidly cycle across generations.

We conclude that E2F activity gradually declines in proportion to the increase in S/G2 length (Fig. 2L). Consequently, cells with short S/G2 phase durations maintain high E2F activity through mitosis, prompting daughter cells to license origins quickly, bypass the restriction point, and continue to cycle. In contrast, when B-MYB/FOXM1 activation is delayed, S/G2 is lengthened, resulting in cells carrying low E2F activity through mitosis, which directs daughter cells to exit to quiescence post-mitosis. Previous work has demonstrated that mitogens are required in S/G2 but not after mitosis in rapidly cycling cells ^2–4^. Thus, high E2F activity sustained through mitosis provides a mechanism through which mitogen sensing in mother cells promotes daughter-cell proliferation.

### Reversal of B-MYB/FOXM1 Activation Triggers G2 Exit and Polyploidy

We next focused on the third cell fate regulated by mitogens in S/G2, the exit of mother cells to polyploid quiescence prior to mitosis (Fig. 1B) ^5^. This exit route represents a special case of an endoreduplication cycle characterized by mitotic bypass combined with CDK2/1 inactivation and cell-cycle exit to quiescence (“G2 exit”) ^36^. G2 exit cells remain in a G1-like state with duplicated DNA, and will enter an additional S phase upon further mitogen stimulation, leading to polyploidy ^5^. Time-lapse analysis in Fig. 1C showed that euploid cells treated with low-mitogen media had greater S/G2 phase duration. We confirmed that a subset of these cells completely inactivates CDK2/1 and exits the cell cycle in G2 before entering mitosis (Fig. 3A, B). In contrast, nearly all cells stimulated with full-serum media progressed rapidly to mitosis (Fig. 3B). Single-cell DNA content analysis using Hoechst staining further confirmed that cells that exited G2 had approximately doubled their DNA content (Fig. 3C), consistent with cells having bypassed mitosis and entered a diploid quiescent state.

**Figure 3.**
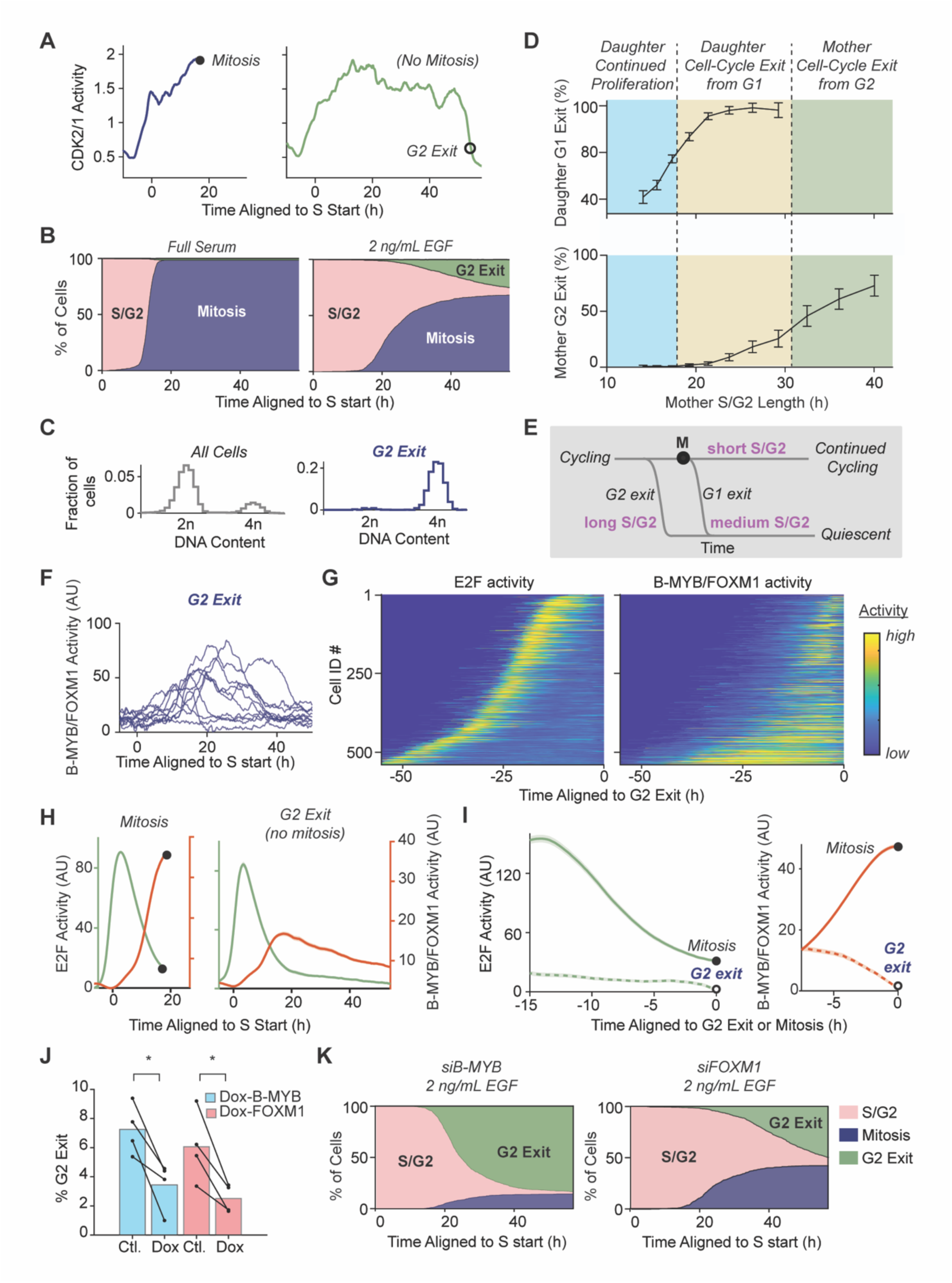
Reversal of B-MYB/FOXM1 Activation Triggers G2 Exit and Polyploidy. **(A)** Left: example of CDK2/1 activity recording from a single cell progressing to mitosis. Right: same for single cell prematurely exiting from G2. **(B)** Cumulative distribution of mitotic entry and G2 exit in cells released with either full-serum media (right, n = 937) or 2 ng/mL EGF (left, n = 1661). **(C)** DNA content histograms derived from Hoechst staining following live-cell imaging, fixed 68 hours after release with 2 ng/mL EGF. Right: DNA content histogram for all cells imaged in condition (n = 15663). Left: DNA content of cells that have exited from S/G2 phase (n = 1291). **(D)** Three ranges of S/G2 length control distinct cell fate outcomes: Short and intermediate S/G2 lengths, daughters enter S or exit G1 (right and middle). Cells exit in G2 with increasing frequency for longer S/G2 durations (left). **(E)** Schematic of the B-MYB/FOXM1 and S/G2 length controlled three-way decision of proliferation cells. **(F)** Representative example time courses of transient B-MYB/FOXM1 increases ending with G2 exit. **(G)** Kymograph of single-cell E2F and B-MYB/FOXM1 time courses in cells ultimately exiting in G2 (2 ng/mL EGF). E2F and B-MYB/FOXM1 activities normalized in single cells to emphasize the relative increases and decreases. **(H)** Mean E2F and B-MYB/FOXM1 activities (+/-SEM), aligned to S start, for cells progressing to mitosis (left, n = 8333 cells) or undergoing G2 exit (right, n = 675 cells) after released with 2 ng/mL EGF. **(I)** Same as (G) but aligned to mitosis or G2 exit (E2F: n = 1391 for mitosis; n = 480 cells for G2 exit); (B-MYB/FOXM1, n = 1146 for mitosis, n = 371 for G2 exit). **(J)** Percent of G2 exit, comparing Control, B-MYB or FOXM1 overexpression. Doxycycline (DOX) or DMSO added 20 h after 2 ng/mL EGF. Only cells after S start. (Four replicates; n = 278 to 3530 cells per condition per replicate; P = .024 and .027 for pcw-MYBL2 and pcw-FOXM1 lines, respectively). **(K)** Cumulative percent mitosis and G2 exit. 2 ng/mL EGF (siB-MYB, n = 1071; siFOXM1, n = 1240).

Since S/G2 is lengthened in cells that exit in G1 or G2, we wondered whether S/G2 length correlates with these cell fates. Intriguingly, three cell outcomes corresponded to different ranges of S/G2 lengths (Fig. 3D): shorter S/G2 lengths direct most daughter cells to continue to proliferate; intermediate lengths direct nearly all daughters to exit to quiescence post-mitosis (standard G1 exit); and only the longest S/G2 lengths were associated with mother cells undergoing mitotic bypass and G2 exit (Fig.3D, E).

Cells monitor the completion of DNA replication (S) at the S/G2 checkpoint, which is thought to initiate mitosis by positive feedback between FOXM1 and CDK2/1, resulting in a sharp increase in CDK2/1 activity ^16,28^. We therefore expected that G2 exit is triggered by failed B-MYB/FOXM1 activation. However, a large majority of cells that exit in G2 did show an increase in their B-MYB/FOXM1 activity (Fig. 3F). Moreover, analysis of both E2F activity and B-MYB/FOXM1 activities in the same cells revealed that while E2F activity was low for periods exceeding 40 hours before the time of G2 exit, B-MYB/FOXM1 remained at intermediate levels prior to G2 exit (Fig. 3G).

To better understand the timing of the E2F and B-MYB/FOXM1 activity changes during G2 exit, we aligned the E2F and B-MYB/FOXM1 activity traces of individual cells to the time of either S phase entry, G2 exit, or mitosis (Fig. 3H, I). When aligned to S start, recordings from cells entering mitosis and those exiting G2 both exhibited indistinguishable average transient increases in E2F activity, as well as initial rises in B-MYB/FOXM1 (comparison of green and red time courses in the left versus right panels, Fig. 3H). However, the B-MYB/FOXM1 time courses started to differ later; cells undergoing G2 exit slowed their activity increase and started to decline their B-MYB/FOXM1 activity. In cells that underwent G2 exit, B-MYB/FOXM1 activity remained elevated for various times before exit (Fig. 3G). When cells were aligned by G2 exit, the averaged B-MYB/FOXM1 activity declined prior to exit, while E2F was already persistently low (Fig. 3I). The corresponding activity traces in cells entering mitosis are included in the two plots as a reference. Together, these results suggest that intermediate B-MYB/FOXM1 rather than E2F activity maintains cells for long periods in G2 and that declining B-MYB/FOXM1 subsequently drives G2 exit.

We next tested the hypothesis that a decrease in B-MYB/FOXM1 activity triggers G2 arrest. If the hypothesis is correct, an increase in relative B-MYB/FOXM1 activity should decrease the proportion of cells that exit in G2. We acutely increased B-MYB/FOXM1 activity by Doxycycline (DOX) induction of B-MYB or FOXM1, both of which lowered the fraction of cells triggering G2 exit (Fig. 3J). Moreover, lowering B-MYB or FOXM1 activity should increase the proportion of cells that exit in G2. Indeed, knocking down B-MYB or FOXM1 greatly increased the percentage of cells exiting in G2 (Fig. 3K, control in Fig. 3B).

We conclude that B-MYB/FOXM1 is not only required for mitosis but also that the B-MYB/FOXM1 program determines how long cells can remain in S/G2. Intermediate B-MYB/FOXM1 activity can keep cells in a G2 maintenance state for long periods and subsequent inactivation of B-MYB/FOXM1 triggers G2 exit.

### Linked B-MYB/FOXM1 and CDK2/1 Inactivation Controls G2 Exit

The positive feedback between B-MYB/FOXM1 and CDK2/1 ^16,28^ raises the question of how B-MYB/FOXM1 activity changes relative to CDK2/1 activity during G2 exit. Single cell analysis demonstrated that cells exiting from G2 increased B-MYB/FOXM1 to varying degrees before inhibiting B-MYB/FOXM1 during G2 exit (Fig. 4A and Fig. S4A). A phase-plot representation of simultaneously recorded B-MYB/FOXM1 versus CDK2/1 activity changes shows that, on average, both activities increase in parallel prior to mitosis (Fig. 4B). Moreover, the activities decrease in parallel in cells exiting from G2. Before cells undergo G2 exit, however, CDK2/1 activity declines on average first, before B-MYB/FOXM1 begins to drop (Fig. 4B), suggesting that for constant mitogen stimulation, CDK2/1 inactivation initiates the G2 exit. Together, these results support that B-MYB/FOXM1 and CDK2/1 activities reinforce each other to first increase and then stay in G2 and suggest that their joint decline drives G2 exit.

**Figure 4.**
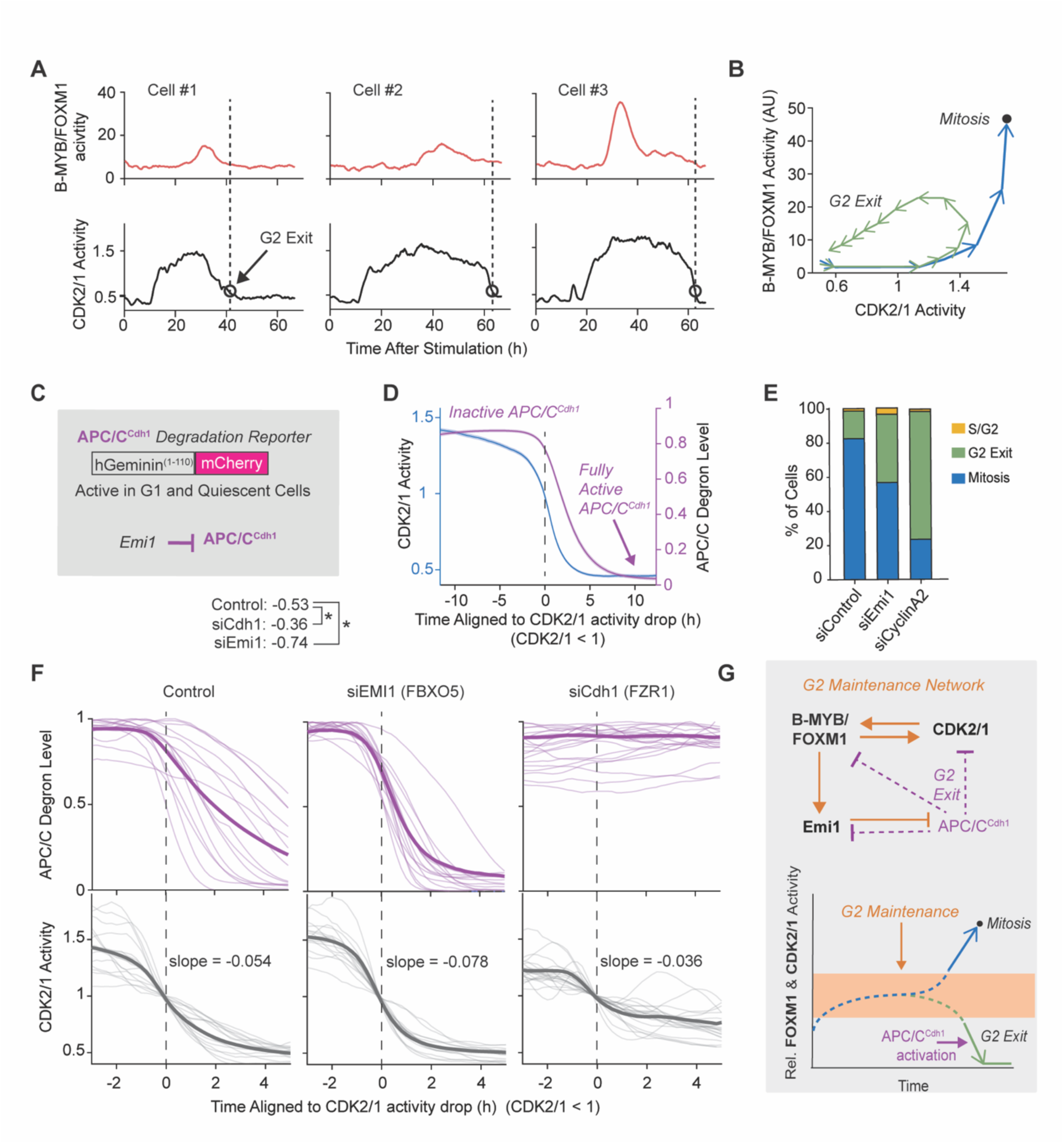
G2 Exit Triggered by Parallel B-MYB/FOXM1 and CDK2/1 inactivation and APC/CCdh1 activation. **(A)** Selected single-cell examples of B-MYB/FOXM1 and CDK2/1 activity changes prior to G2 exit, emphasizing that B-MYB/FOXM1 activities can increase to high levels and still reverse. **(B)** Averaged phase plot of single-cell B-MYB/FOXM1 versus CDK2/1 activities (n = 1007, +/-SEM). 2 ng/mL EGF. Arrows represent 4 h; 32 h period before G2 exit. **(C)** Schematic; APC/C^Cdh1^ degron reporter and APC/C^Cdh1^ inhibitors and targets. **(D)** Mean traces of CDK2/1 activity versus APC/C^Cdh1^ reporter degradation during G2 exit. Aligned by CDK2/1 < 1; 2 ng/mL EGF (n = 480). **(E)** Percent G2 exit, S/G2 and mitosis cell fates. Comparing siControl, siEMI1, and siCCNA2. 72 h after 2 ng/mL EGF. **(F)** Single-cell and average CDK2/1 and APC/C^Cdh1^ reporter activities in cells exiting G2. Aligned to CDK2/1 activity drop below 1. 2 ng/mL EGF. Comparison: siControl, siFZR1, or siEMI1. n = 31 - 130 cells per condition. CDK2/1 slope quantified by linear regression; two-tailed t-test (p = 1.36*10^−8^ for siEMI1 vs. siCTRL; p = 3.30 * 10^−3^ for siFZR1 vs. siCTRL). Dashed line, approximate amplitude for inactive CDK2/1. **(G)** Schematic of the regulation of the G2 maintenance network (top) from which cells can enter mitosis or undergo G2 exit (bottom).

An additional cell cycle regulator implicated in endoreduplication is the E3 ligase APC/C^Cdh1^. In a healthy cell cycle, APC/C^Cdh1^ is thought to be exclusively active in G1 and in G0, where it degrades substrates such as Cyclin A2, FOXM1, E2F7/8, and EMI1 (encoded by gene FBXO5) (Fig. 4C) ^5,7,37–39^. Thus, activation of APC/C^Cdh1^ is thought to be a general hallmark of exit to quiescence (Fig. S3A). Prior work suggests that E2F and FOXM1 can keep APC/C^Cdh1^ inhibited by inducing the APC/C^Cdh1^ inhibitor Emi1 ^22^ and APC/C^Cdh1^ can be further inhibited by phosphorylation of Cdh1 by CDK2/1 ^21^. To understand the role of APC/C^Cdh1^ in G2 exit, we used a dual CDK2/1 and APC/C^Cdh1^ reporter cell line (Fig. 4D) ^21^. Prior to G2 exit, APC/C^Cdh1^ was initially inactive when CDK2/1 activity began to drop but became reactivated as CDK2/1 activity continued to decline (Fig. 4D; single-cell examples in Fig. S3B).

We found that premature activation of APC/C^Cdh1^ promotes G2 exit, as demonstrated by the fact that knockdown of the APC/C^Cdh1^ inhibitor Emi1 ^37^ increased the percentage of cells exiting in G2 (Fig. 4E; single-cell examples in Fig. S3C). As a positive control, the effect of cyclin A2 knockdown on G2 exit is also shown (Fig. 4E). To understand the role of APC/C^Cdh1^ in promoting G2 exit, we examined the effect of APC/C^Cdh1^ activity on the kinetics of CDK2/1 inactivation during G2 exit (Fig. 4F, right). Emi1 knockdown accelerated the rate of the decline in CDK2/1 (Fig. 4F, middle). Next we tested the effect of knockdown of Cdh1 (encoded by FZR1) ^21^, which keeps APC/C^Cdh1^ inactive. Markedly, CDK2/1 activity still declined with inactive APC/C^Cdh1^, but more slowly and to a CDK2/1 activity above basal (Fig. 4F, left). Thus, APC/C^Cdh1^ functions as an accelerator of CDK2/1 inactivation that is also needed to fully inactivate CDK2/1 and complete G2 exit.

Our results suggest that G2 exit is primarily controlled by declining B-MYB/FOXM1, and not E2F, even though both activities can keep CDK2/1 active. We observe that cells do not undergo G2 exit until E2F activity reaches very low levels at ∼20 hours after S-phase entry, suggesting that E2F activation may protect cells from G2 exit. However, once E2F becomes inactive, B-MYB/FOXM1 activity in turn prevents G2 exit. In mitogen-limited conditions, cells may remain for tens of hours in G2 phase before committing to either mitosis or G2 exit. These cells enter a G2-maintenance state characterized by intermediate B-MYB/FOXM1 and CDK2/1 activities, requiring expression of the B-MYB/FOXM1 targets Emi1 and cyclin A2 (Fig. 4E and 4G, top) ^22^. Cells can transition out of this G2 maintenance state to enter mitosis (Fig. 4G, bottom), but with increasing time spent arrested in G2, an increasing number of cells trigger G2 exit, initiated by declining CDK2/1 and B-MYB/FOXM1 activity and completed by APC/C^Cdh1^ activation (Fig. 4G, bottom).

## DISCUSSION

The classic cell-cycle restriction point model assumed that E2F is fully inactivated in cycling cells before mitosis, placing the mitogen sensitivity and decision in G1 ^1,8,23^. Here we identify a second B-MYB/FOXM1-regulated restriction point that decides the cell-cycle fate of cycling cells in G2 rather than G1.

Instead of reaching mitosis without E2F activity, rapid B-MYB/FOXM1 activation shortens S/G2, resulting in high E2F activity being carried through mitosis to trigger continued cycling. The high E2F activity bypasses the need for mitogens in G1, allowing cells to keep cycling unchecked (Fig. 5A). Only in cells where B-MYB/FOXM1 activation is delayed and S/G2 lengthened, is mitotic E2F activity low, and cells either temporarily or permanently exit the cell cycle in G1. Such cells either stay quiescent, or they must again activate E2F and pass the classic restriction point to enter the next cell cycle. Thus, B-MYB/FOXM1 controls daughter fates indirectly via the mother’s S/G2 duration, which sets the mitotic E2F activity that determines whether cells continue to cycle or exit in G1. Moreover, if S/G2 extends beyond ∼20 hours, E2F activity becomes inactivated in mother cells. By that time, cells have either already activated B-MYB/FOXM1 enough to enter mitosis or have settled into a G2 maintenance state with intermediate B-MYB/FOXM1 activity. In this maintenance state, cells can inactivate B-MYB/FOXM1, triggering mitotic bypass and exit into a diploid quiescent state.

**Figure 5.**
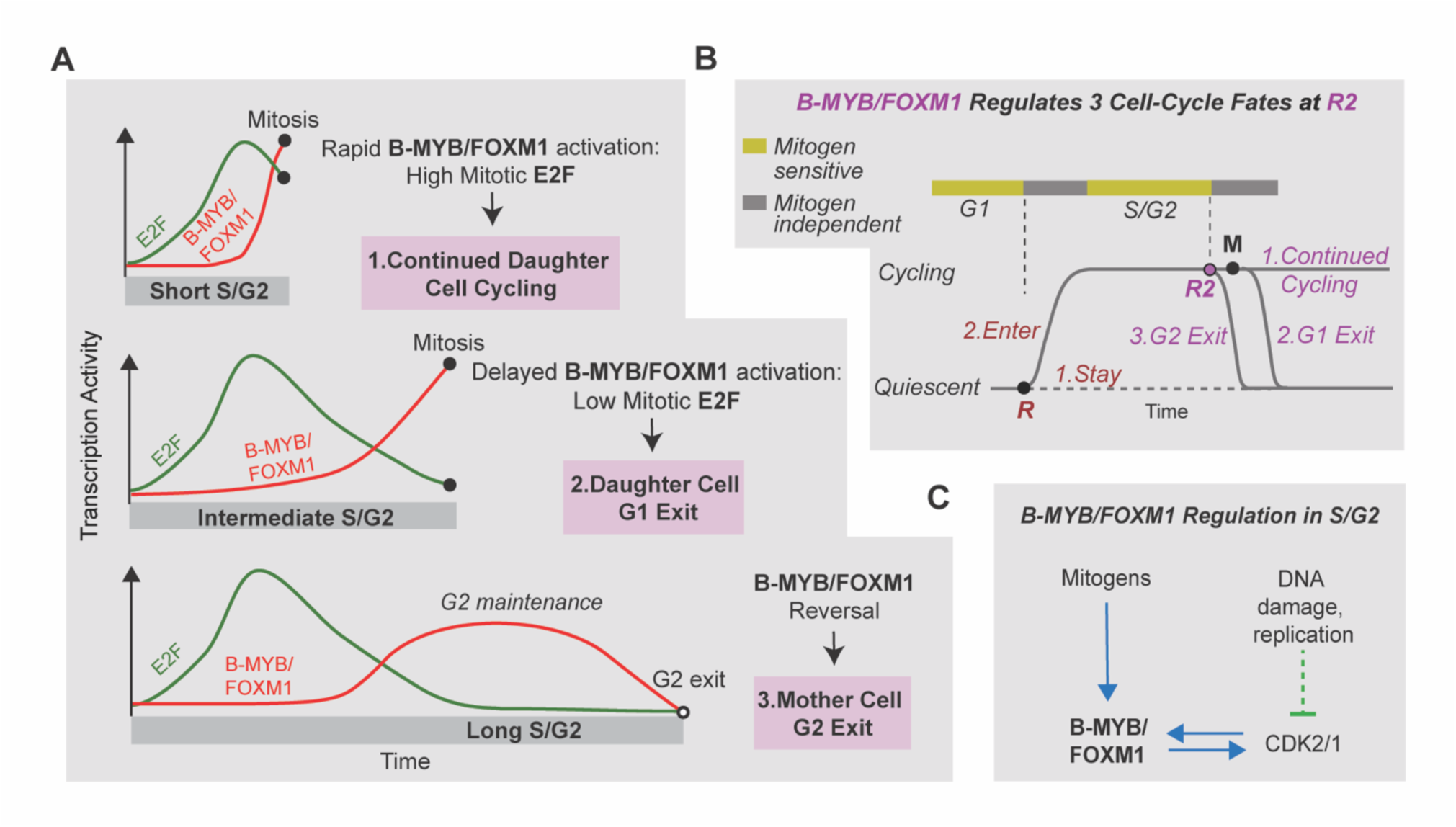
Central Role of B-MYB/FOXM1 in the G2 restriction Point Model. **(A)** Schematic illustration showing how regulation of B-MYB/FOXM1 activation controls S/G2 length, which decides between 3 cell-cycle fates: continued proliferation, standard G1 exit and premature G2 exit. **(B)** Schematic of the G2 restriction point controlling three cell-cycle fates. **(C)** Schematic of the regulation of B-MYB/FOXM1 in S/G2.

Thus, a revised cell-cycle model includes two restriction points that separately control whether cells enter the cell cycle or continue cycling: (1) in quiescent cells, a bi-directional, mitogen-regulated “G1 restriction point” (R), centered on E2F, determines whether cells enter the cell cycle or stay in quiescence (Fig. 5B). (2) In cycling cells, a tri-directional, mitogen-regulated “G2 restriction point” (R2), centered on B-MYB/FOXM1, governs three fates: continued daughter-cell cycling, daughter-cell G1 exit, or mother-cell G2 exit into polyploid quiescence (Fig. 5B).

G2 exit combines mitotic bypass with entry into a diploid quiescent state. Intermediate B-MYB/FOXM1 activity prevents G2 exit by maintaining the expression of its substrates, cyclin A2 and Emi1 (Fig. 4E, G). G2 exit is the result of a coordinated inactivation of B-MYB/FOXM1 and CDK2/1 and activation of APC/C^Cdh1^ that unfolds over several hours. In mitogen-limited conditions, fewer cells proceed into mitosis as S/G2 lengthens, whereas cells undergo G2 exit at a slow but largely constant rate (Fig. 3B). This behavior can explain the previously described cell-cycle timer model, in which cells increasingly exit G2 as S/G2 lengthens ^5^. However, while our data confirms this finding, it also suggests that G2 exit is better described as a regulated process where changes of positive mitogen signal increase B-MYB/FOXM1 activity to trigger mitosis or changes of inhibitory signal inhibit B-MYB/FOXM1 to trigger G2 exit (Fig. 3H, 4G).

Our findings support that B-MYB/FOXM1 activity is regulated in S/G2 through diverse opposing mitogen and stress signaling mechanisms (Fig. 5C). B-MYB and FOXM1 expression are primarily regulated by E2F ^22^; FOXM1 is also induced by autoregulation ^40^, MYC ^41^, YAP/TEAD ^42^, and by B-MYB ^43^. However, expression alone is not sufficient: B-MYB and FOXM1 must be phosphorylated to become active. While CDK2/1 is the main kinase that phosphorylates and activates B-MYB/FOXM1 (Fig. 1G–I), CDK4/6 is also known to phosphorylate FOXM1 ^44^. In addition, stress signals regulate B-MYB/FOXM1: DNA damage and replication stress suppress CDC25, thereby reducing CDK2/1 activity ^32,45,46^, which indirectly lowers FOXM1 activity ^16^. DNA damage can also activate p53 (TP53), which expresses p21 to inactivate CDK2/1 ^47^.

Cells that exit in G2 are recognized by a lack of proliferation markers and diploid/polyploidy DNA content. Based on such histology analysis, B-MYB/FOXM1-regulated G2 exit may be particularly important in specialized mammalian tissues that are frequently polyploid, such as the liver ^48^ and pancreatic β-cells ^49^, as well as in many cancers ^50^. This is consistent with *in vivo* observations that polyploid quiescence is rare in humans and rodents ^51^. By contrast, B-MYB/FOXM1 and the G2 restriction point (R2) may have a general role in cycling cells, determining when cells undergo intergenerational cycling for several generations and when to return to quiescence.

Our study also has therapeutic implications. B-MYB and FOXM1 overexpression is found across many cancers and correlates with poor outcomes ^11,52,53^, yet the mechanistic link to aggressive proliferation has been unclear. Our results suggest that elevated B-MYB and FOXM1 helps cancer cells keep cycling across generations despite the heightened DNA damage common in cancer ^54^ that would otherwise trigger G1 or G2 exits. This role of B-MYB/FOXM1 provides mechanistic support for B-MYB/FOXM1-targeted therapies ^55–58^. In conclusion, we show that B-MYB/FOXM1 regulation in G2 controls three cell-cycle fates: continued cycling, standard G1 quiescence, and premature diploid quiescence, opening new therapeutic opportunities.

**Table 1:**
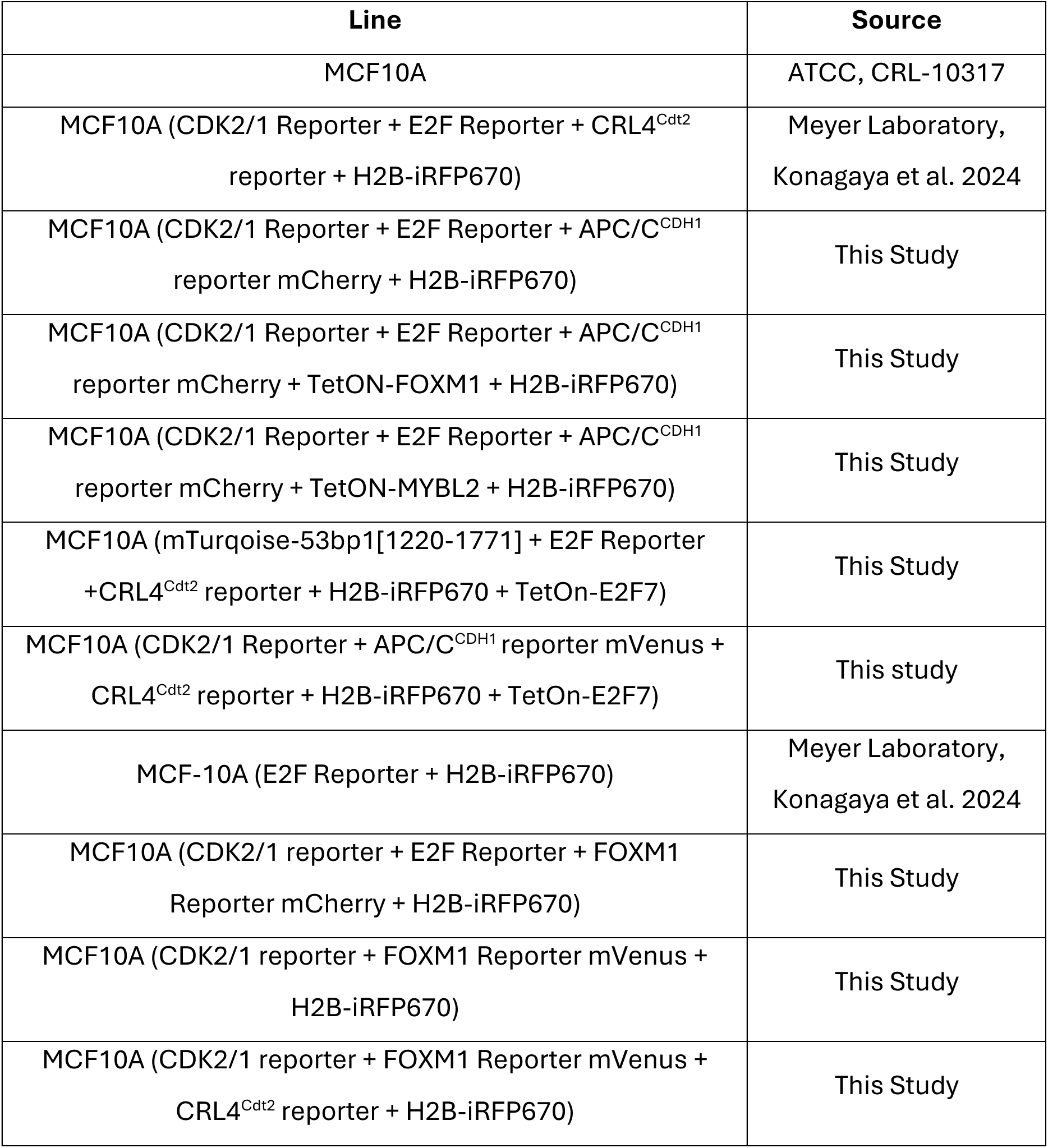
Cell lines used in this study.

---

**Figure S1.**
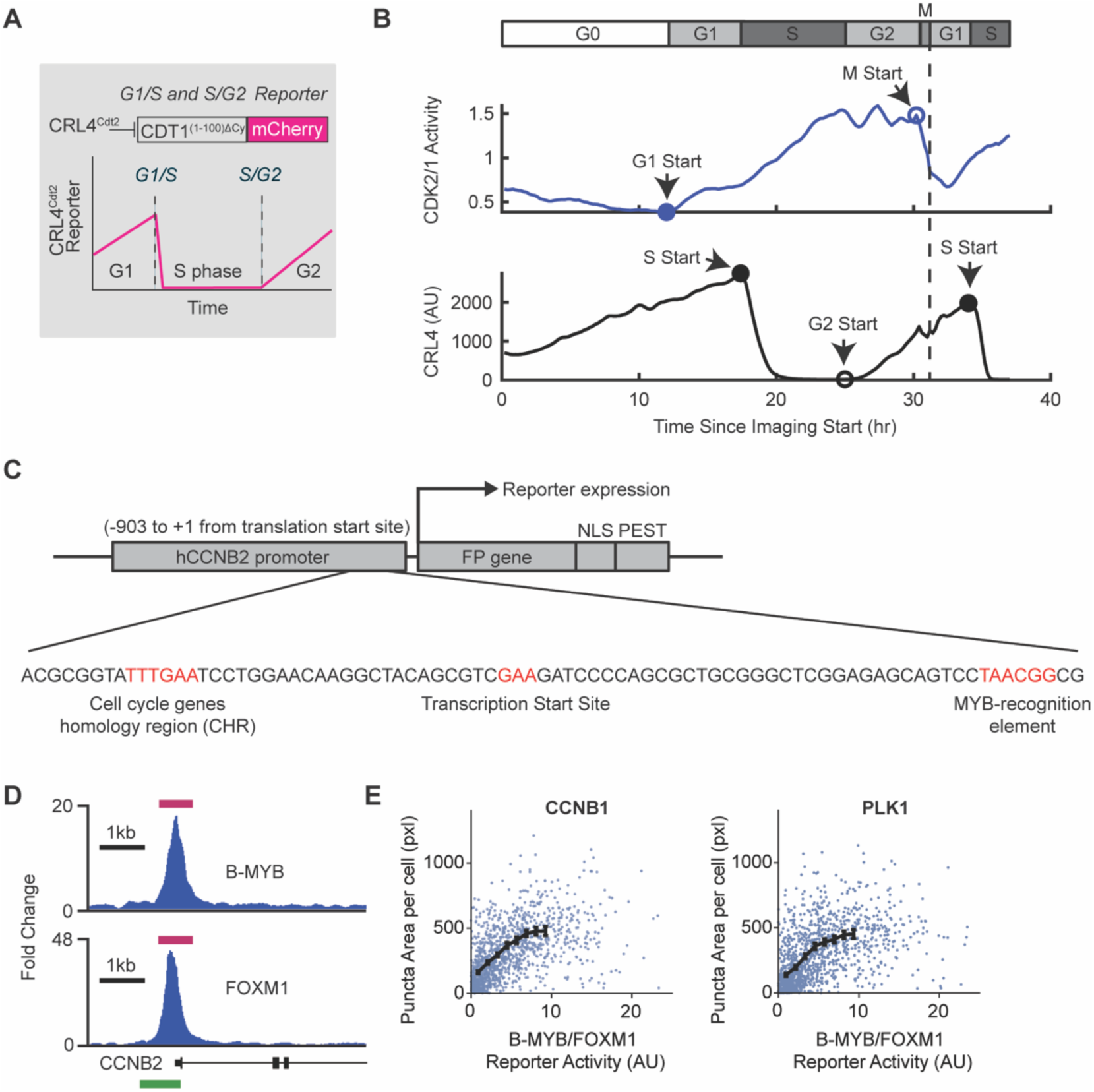
**(A)** Schematic of FUCCI(CA) CRL4^Cdt2^ reporter to measure the G1/S and S/G2 transitions. **(B)** Single-cell traces of the CDK2/1 and CRL4^Cdt2^ reporter activities with marked cell cycle phase transitions. **(C)** Design of the B-MYB/FOXM1 transcriptional reporter system. *hCCNB2* promoter region was used for the reporter development. *hCCNB2* promoter has both cell cycle genes homology region (CHR) and MYB-recognition element. **(D)** B-MYB and FOXM1 ChIP-Seq reads (blue) over *hCCNB2* promoter region used in the B-MYB/FOXM1 reporter (green box). Magenta lines indicate detected peaks with a discovery rate (IDR) cutoff = 0.05 (from ENCODE database). **(E)** Single-cell correlation of B-MYB/FOXM1 reporter intensity against mRNA FISH puncta area for *CCNB1* (left, r = .63, p < 10^−308^) or *PLK1* (right, r = .61, p < 10^−308^), respectively. N = 2000 cells per scatter plot. Trend line shows the mean of binned data +/-2*SEM based. Statistics based on 4217 (CCNB1) or 3781 (PLK1) cells. Cells were fixed 25 h after release with 2 ng/mL EGF.

**Figure S2.**
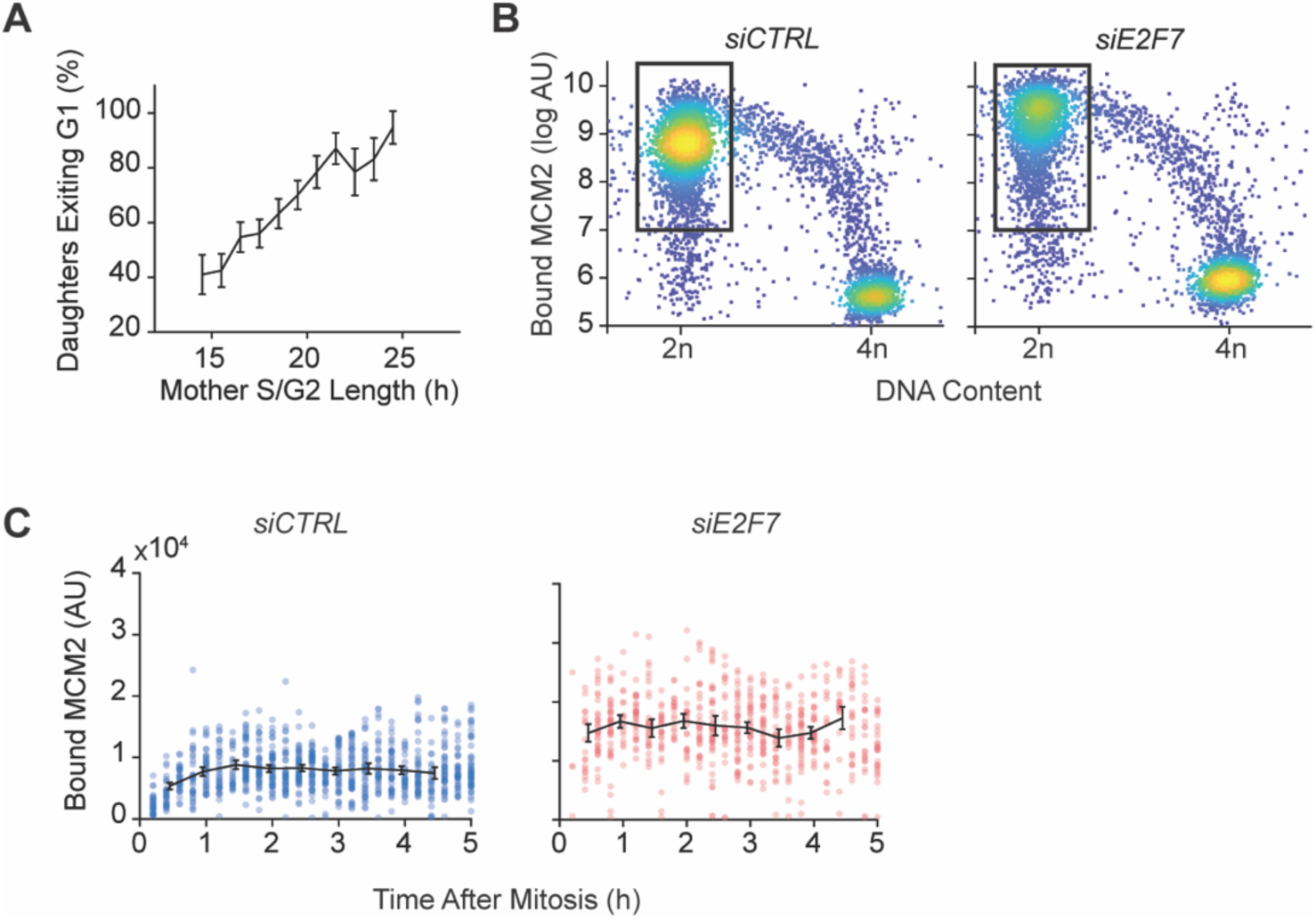
**(A)** Percent of daughter cells undergoing G1 exit versus their mothers S/G2 length (n = 2724), binned by S/G2 length. Error bars, 2*SEM. **(B)** Chromatin-bound MCM2 versus DNA content, same conditions as Fig. 2h (siCTRL, n = 4500 cells per plot). Rectangles highlight the cells plotted in Figure 2I. **(C)** Time-course analysis of chromatin-bound MCM2 plotted over time aligned by the end of mitosis (siCTRL, n = 1058; siE2F7, n = 691; see Methods).

**Figure S3.**
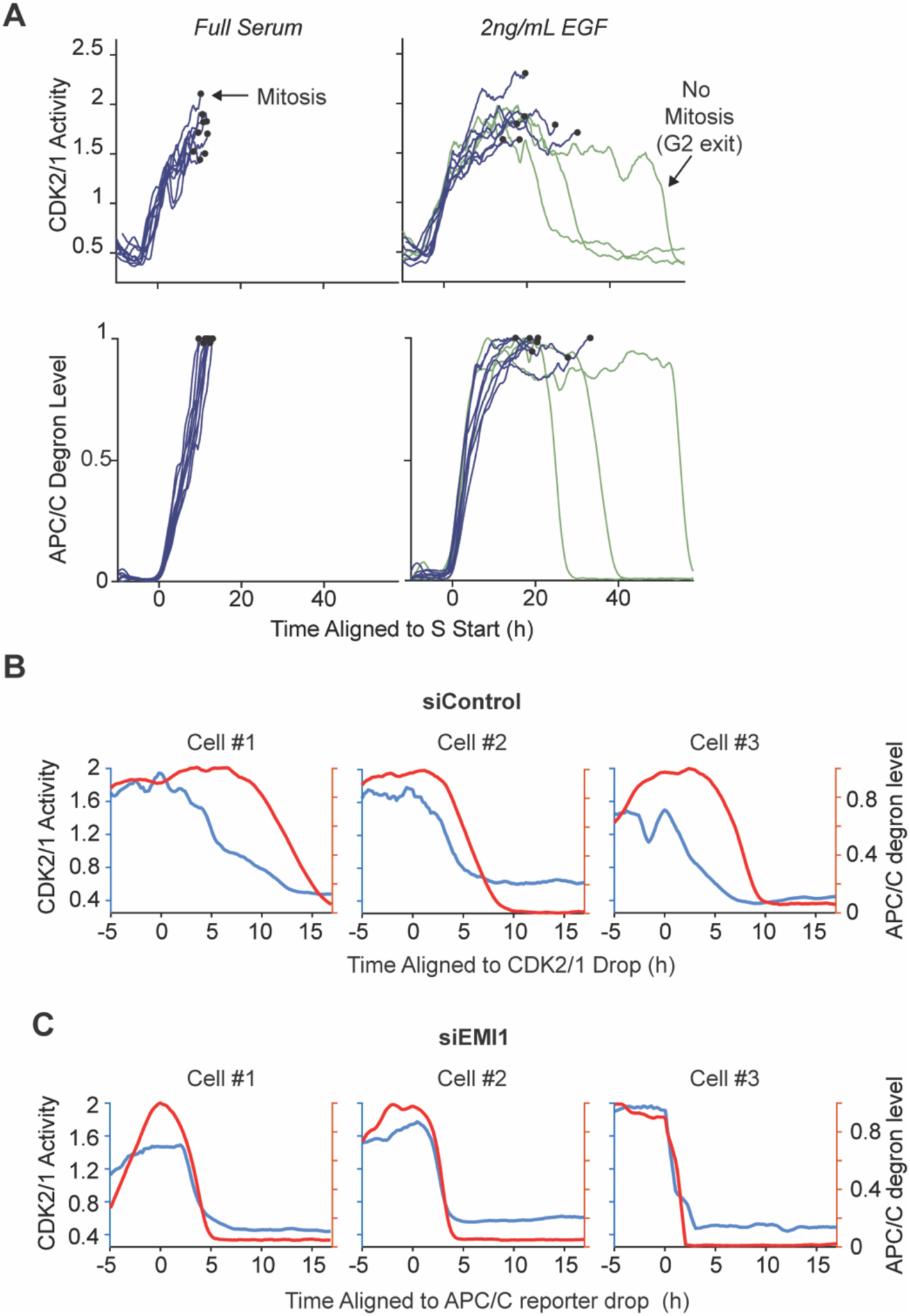
**(A)** Examples of CDK2/1 and APC/C^Cdh1^ activity traces of cells entering mitosis or undergoing G2 exit; full serum versus 2 ng/mL EGF. **(B)** Single cell comparisons of the decline in CDK2/1 activity and increase in APC/C^Cdh1^ activity. CDK2/1 inactivation generally precedes the activation of APC/C^Cdh1^. **(C)** siEmi1 application accelerates the CDK2/1 declines in parallel with the increase in APC/C^Cdh1^ activity.

## METHODS

### Cell Culture

MCF10A human mammary epithelial cells (sourced from ATCC) were utilized for all experiments unless specified otherwise. These cells were maintained in DMEM/F12 growth medium containing HEPES (Gibco, 11039047) with the addition of 5% horse serum (Gibco, 16050122), 20 ng/mL EGF (PeproTech, AF-100-15), 0.5 μg/mL hydrocortisone (Sigma, H0888), 100 ng/mL cholera toxin (Sigma, C8052), and 10 μg/mLinsulin (Sigma, I1882). For cell passage, trypsin-EDTA solution (0.05%, Gibco, 25300054) was employed, followed by neutralization using DMEM/F12 containing 20% horse serum. Lenti-X 293T human embryonic kidney cells (Takara Bio, 632180, RRID: CVCL_4401) were grown in DMEM supplemented with 10% FBS. For serum starvation of MCF-10A cells, growth medium lacking horse serum, EGF, and insulin but containing 0.3% BSA was used after washing cells three with this starvation medium. Mitogen release was achieved by replacing starvation medium with either starvation medium supplemented with EGF or complete growth medium. All cell cultures were maintained at 37°C with 5% CO2. For imaging experiments, 96-well glass-bottomed plates (Cellvis, P96-1.5H-N) were prepared by coating with collagen (Advanced Biomatrix, 5005-B, 30–60 μg/mL for minimum 20 minutes). Cells were plated in these wells at least one night prior to experimental procedures.

### Cell Line Generation

Constructs were introduced into cells using second- or third-generation lentiviral transduction. For second-generation transduction, HEK-293T cells were co-transfected with packaging plasmids psPAX2 (Addgene #12260) and pMD2.G (Addgene #12259) and the lentiviral plasmid together with Lipofectamine 2000 (Thermo, 11668019). For third-generation transduction, packaging plasmids pMDLg/pRRE (Addgene #12251), pRSV-rev (Addgene #12253) and pCMV-VSV-G (Addgene #8454) were used. Viral particles were harvested from supernatant 72 hours post-transfection, passed through a 0.22 μm filter (Millipore, SCGP00525), and concentrated using 100 kDa centrifugal filters (Millipore, UFC910024). The concentrated virus was subsequently used to transduce target cells in growth medium. For constitutively expressed fluorescent constructs lacking antibiotic resistance selection markers, cells showing positive fluorescence were isolated using a BD Aria II cell sorter (at the Weill Cornell Medicine Shared FACS Facility). For cells expressing constructs with puromycin selection markers, selection was performed using 1 μg/mL puromycin until control cells were no longer viable. These cell lines were maintained without doxycycline until experimental use. To establish clonal cell lines expressing the B-MYB/FOXM1 reporter, mVenus-positive MCF10A cells underwent single-cell cloning. MCF10A cells were obtained directly from ATCC, while Lenti-X 293T cells were sourced directly from Takara Bio. All cell lines were confirmed negative for mycoplasma contamination. See Table 2.1 for a complete accounting of cell lines used in this study.

### Plasmid Generation

The plasmids created during this research were constructed through Gibson assembly of PCR-amplified inserts and plasmid backbones, cut via restriction enzyme digestion. To generate the B-MYB/FOXM1 reporter, the *CCNB2* promoter region (nt −903 to +1 relative to translational start) was amplified out of purified genomic DNA from RPE-1 cells and then inserted into the backbone of pLV-hCDC6p-Venus (Addgene #212665). To generate the Dox-inducible constructs pCW-MYBL2-HA-puro and pCW-FOXM1-HA-puro, human MYBL2 and FOXM1 cDNA was obtained as synthesized gene fragments and inserted into the pCW backbone (derived from pCW-Cas9, Addgene #50661). The B-MYB/FOXM1 reporter and other constructs used in the study are available through the non-profit organization Addgene (https://www.addgene.org).

### siRNA Transfection

MCF10A cells were transfected using DharmaFECT 1 reagent (Dharmacon, T-2001-03) following the manufacturer’s recommended protocol. The transfection mixture contained 2 nM siRNA combined with DharmaFECT 1 at a final dilution of 1:500. The cells were incubated in transfection mixture for 5 to 24 hours in either growth medium or serum-starvation medium, after which the medium was changed. For live-cell imaging experiments, cells were transfected either immediately prior to, or overnight the night before, imaging start. Pools of four siRNA oligonucleotides (ON-TARGETplus, Dharmacon) were used for siControl, siMYBL2, siFOXM1, siFBXO5, siCCNA2, and siE2F7.

### Chemicals

The following stock solutions of drugs were dissolved in DMSO (Sigma, D2650 or Santa Cruz, sc-358801): doxycycline hyclate (Sigma, D9891); the CDK2 inhibitor PF-07104091 (ChemieTek, CT-PF0710); the CDK1 inhibitor RO-3306 (Cayman Chemical, 15149); the Wee1 inhibitor MK1775 (Cayman Chemical, 21266).

### RNA Fluorescence In-Situ Hybridization

Cells were fixed with 4% paraformaldehyde in PBS for 10 minutes at room temperature then washed with PBS. Cells were next permeabilized with 0.2% Triton X-100 in PBS for 15 minutes, with a subsequent PBS wash. RNA fluorescent in situ hybridization (FISH) was conducted using the ViewRNA ISH cell assay (Thermo, QVC0001) in accordance with the manufacturer’s protocol. Following FISH, cells were washed with PBS and stained with 1 μg/mL Hoechst 33342 (Invitrogen, H3570) in PBS for 15 minutes then PBS-washed again prior imaging. The hybridization probes employed in this study were: CCNB1 (VA6-16942-VC) and PLK1 (VA6-3167940-VC).

### Immunofluorescence

#### General Protocol

Cells were fixed with 4% paraformaldehyde in PBS for 15 minutes at room temperature, followed by a PBS wash. If cells expressed fluorescent proteins with spectral overlap with subsequently used fluorophores, these proteins were chemically bleached ^60^ using a solution of 3% H2O2 with 20 mM HCl in PBS for 1 hour, followed by PBS wash. Cells were permeabilized in 0.2% Triton X-100 in PBS for 15 minutes then blocked for 1 hour in buffer containing 10% FBS, 1% BSA, 0.1% Triton X-100, and 0.01% NaN3 in PBS. Primary antibody incubation occurred overnight in blocking buffer at 4°C, followed by a PBS wash and secondary antibody application in blocking buffer for 1 hour at room temperature. After a PBS wash, nuclear counterstaining was performed using 1 μg/mL Hoechst 33342 (Invitrogen, H3570) in PBS for 15 minutes, with a final PBS wash before imaging. PBS washes were performed either with an automated plate washer (BioTek 405 LS) using a protocol of aspiration to 50 μL followed by dispensing 250 μL, repeated 3 times, or manually (specifically for pre-extracted samples, involving 3 complete aspiration cycles).

#### Pre-extraction for chromatin-bound protein

For chromatin-bound protein detection, soluble proteins were extracted prior to fixation. Immediately after culture medium removal, plates were placed on ice. Cells underwent a 5-minute incubation in ice-cold pre-extraction buffer containing 0.2% Triton X-100 (Sigma, X100) and 1× Halt Protease Inhibitor cocktail (Thermo, 78429) in PBS. Following pre-extraction, 8% paraformaldehyde in H2O was added directly to the wells in a 1:1 ratio using wide-orifice tips to minimize cellular detachment. Fixation proceeded for 30 minutes at 4°C, after which staining proceeded according to the standard immunofluorescence protocol.

### Antibodies

The following primary antibody was used: rabbit anti-MCM2 (1:800, Cell Signaling Technology 3619). Secondary antibodies targeting the appropriate species and with no spectral overlap were selected from the following (diluted 1:1000): goat anti-mouse IgG Alexa Fluor 647 (Thermo A-21235), goat anti-mouse IgG Alexa Fluor 514 (Thermo A-31555).

### Microscopy

Automated epifluorescence microscopy was conducted using a Ti2-E inverted microscope (Nikon) equipped with an 89903-ET-BV421/BV480/AF488/AF568/AF647 Quinta Band filter set (Chroma Technology), illuminated by an LED light source (Lumencor Spectra X). Images were captured through a 20x objective (Nikon CFI Plan Apo Lambda, 0.75 NA) via an ORCA-Flash4.0 V3 sCMOS camera (Hamamatsu). All images were collected in 16-bit format with 2 × 2 binning, with exposure parameters selected to prevent signal saturation. For time-lapse experiments with living cells, 96-well plates containing 200 μL of medium per well were monitored within an environmental chamber maintained at 37°C with 5% CO2. Each experimental condition was represented by at least two wells, with 9 distinct sites imaged per well at 12-minute intervals. To minimize phototoxicity, exposure times were kept to the minimum required for adequate signal-to-noise ratio for each channel, with total light exposure restricted to under 100 ms per site at each time point. In protocols involving live-cell imaging followed by fixed-cell analysis, specimens were promptly removed from the microscope after the final time point and immediately fixed. When correlating fixed-cell images with either previous live-cell data or earlier rounds of fixed-cell imaging, the plate position (which may shift when the plate is repositioned on the microscope) was manually aligned to approximately match the original location, with precise registration subsequently achieved through computational alignment during image analysis.

### Cell-Cycle Reporters

Cell cycle reporters of CRL^Cdt2^, APC/C^Cdh1^ (FUCCI geminin) and CDK2/1 activity were used in this study. The CRL4^Cdt2^ and APC/C^Cdh1^ reporters were derived from the previously established FUCCI(CA) reporter system. The CRL4^Cdt2^ activity reporter consists of a human CDT1 fragment (amino acids 1-100), designated as CDT1(1-100)ΔCy in the FUCCI(CA) system, which lacks origin licensing functionality. This construct incorporates a PCNA-interacting protein (PIP) degron that facilitates CRL4^Cdt2^-mediated degradation when PCNA associates with activated origins. Additionally, the Cy motif was eliminated to prevent SCF^Skp2^-dependent degradation. This reporter exhibits characteristic dynamics, with rapid degradation to minimal levels at S-phase initiation and subsequent re-accumulation at G2 phase onset. The APC/C^Cdh1^ reporter, corresponding to Geminin(1-110) in the FUCCI(CA) system, comprises the first 110 amino acids of human Geminin fused to mCherry. Geminin(1-110) undergoes degradation during anaphase via APC/C^Cdc^^20^ and throughout G1 via APC/C^Cdh1^, followed by re-accumulation when APC/C^Cdh1^ is inactivated at the G1/S transition. While these reporters are commonly assessed by presence/absence in single-timepoint analyses, time-lapse microscopy of reporter fluorescence dynamics in individual cells enables precise determination of CRL4^Cdt2^ activation and inactivation (marked by the initiation of CRL4^Cdt2^ reporter degradation or re-accumulation, respectively) and APC/C^Cdh1^ inactivation or reactivation (indicated by geminin reporter stabilization or degradation).

The CDK2/1 activity reporter functions through a translocation mechanism, undergoing phosphorylation by complexes of cyclin E or A with CDK2 or CDK1. This construct is based on a fragment of human DNA helicase B (residues 994-1087) that serves as a substrate for cyclin E/A-CDK2/1 phosphorylation ^4^. During G0 and early G1 phases, the unphosphorylated reporter localizes to the nucleus, but progressively translocates to the cytoplasm as cyclin E/A-CDK2/1 activity increases throughout the cell cycle, due to enhanced phosphorylation.

Thus, the cytoplasmic to nuclear intensity ratio reflects instantaneous cyclin E/A-CDK2/1 activity. Crucially, CDK2/1 activity must reach a threshold for S-phase entry of approximately 0.7-1, depending on media conditions. For this study a CDK2/1 activity threshold of 0.9 was used to approximate the start of S-phase entry.

### Image Processing and Single-Cell Tracking

Image processing was conducted using a customized Matlab pipeline as previously documented ^21,61^. The procedure involved initial flatfield correction (illumination bias was determined by aggregating background regions from multiple wells within the same imaging session), followed by local background subtraction. Nuclear segmentation was based on either Hoechst staining or H2B signal. For analyses involving E2F, B-MYB/FOXM1, Geminin, and CRL4^Cdt2^ reporters, the median signal intensity within each nucleus was quantified. CDK2 activity was determined by calculating the ratio of median cytoplasmic intensity to median nuclear intensity. The cytoplasmic region was defined by creating an expanded ring outside the nucleus (with inner radius set at 0.65 μm and outer radius at 3.25 μm), ensuring no overlap with neighboring cells. For RNA FISH quantification, whole-cell regions were segmented by approximating an area containing the nucleus and extending up to 15.6 μm beyond the nuclear boundary without overlap of adjacent cells. FISH puncta were identified by applying a top-hat filter to the raw images using a circular kernel with 1.3 μm radius, followed by absolute intensity thresholding. RNA puncta count was determined as the number of foreground pixels within each defined whole-cell region. The complete image analysis pipeline for this study will be deposited in GitHub (https://github.com/MeyerLab).

### Cell-cycle Annotation of Live-cell Data

S-phase start was annotated (depending on the reporter’s combinations available in each cell line) by either the CRL4^Cdt2^, APC/C^Cdh1^, or CDK2/1 reporters, as indicated in figure legends. We identified CRL4^Cdt2^ activation (characterized by the initiation of CRL4^Cdt2^ reporter degradation) and Geminin degradation onset by applying a drop detection algorithm to traces following mitosis or mitogen stimulation. This algorithm identifies reporter degradation at specific timepoints based on a predetermined number of subsequent frames (typically ∼3 frames, which represents the minimum detectable interval since degradation initiation). By using a fixed number of frames beyond the degradation point for detection, we eliminated potential biases in accuracy between cells exhibiting recent degradation versus those with earlier degradation events. For APC/C^Cdh1^ inactivation (defined as the start of APC/C^Cdh1^ reporter accumulation), we employed a complementary detection approach, pinpointing the initial timepoint where the slope increases to a threshold value and the reporter transitions from low levels to sustained higher levels. G2 start was similarly identified as the initial timepoint at which the CRL4^Cdt2^ reporter begins to accumulate after remaining low in S-phase, with the slope of the reporter signal increasing above a threshold value. In the absence of the CRL4^Cdt2^ reporter, G2 phase was approximated, in 2ng/mL EGF media, as 8 hours after S-phase start as per the CDK2/1 reporter.

G2 dropout was identified via the CDK2/1 reporter as the first point at which the CDK2/1 activity begins to persistently drop before remaining below a threshold of CDK2/1 activity < 0.6, without completing cell division. For G2 dropout analysis, analysis was limited to cells that either divided or were tracked for the entire analyzed imaging duration. Mitosis was annotated as part of the cell tracking algorithm, with mitosis defined as the timepoint at which the two sets of chromosomes separate during anaphase. All threshold parameters were established empirically and visually confirmed through examination of a minimum of 200 individual traces.

### ChIP-seq Analysis

B-MYB and FOXM1 ChIP-seq data, including signals and peaks associated with the *CCNB2* promoter region, were obtained from the ENCODE database (https://www.encodeproject.org/ ^62^); accession numbers: ENCFF762JVU.bigWig, ENCFF739GSE.bigBed, ENCFF275EAX.bigWig, ENCFF739GSE.bigBed; cell line: K562. All experimental procedures were conducted in duplicate, with an irreproducible discovery rate threshold of 0.05.

### Statistical Analysis

All experiments were repeated at least 2 times on different days unless noted. Each condition was evaluated in 2-4 different wells in 96-well plates with time-courses and single-cell reporter activities from several hundred cells per well individually analyzed. Statistical evaluations were conducted using two-sided, two-sample t-tests. Results are presented as either mean ± standard error or median with 25th and 75th percentiles, as indicated in figure legends. Statistical significance is denoted as follows: \**P* < 0.05, \*\**P* < 0.01 and \*\*\**P* < 0.001. Correlations of fixed stains to E2F activity (Fig. 2I) were calculated via Spearman’s rank correlation coefficient, while linear fits were derived from least square regression. The correlation of E2F activity with S/G2 length (Fig. 2C) was fit to a two-term exponential model using robust fitting with a bisquare weight function. Additional statistical parameters for experiments are detailed within respective figure legends. Sample size was not statistically determined a priori. Randomization was not implemented in the experimental design, and researchers were not blinded during experimental procedures or when assessing outcomes.

